# Brain Iron Imaging Markers in the Presence of White Matter Hyperintensities

**DOI:** 10.1101/2021.03.22.436449

**Authors:** Kyle D. Murray, Madalina E. Tivarus, Giovanni Schifitto, Md Nasir Uddin, Jianhui Zhong

## Abstract

**Purpose:** To investigate the relationship between pathological brain iron deposition and white matter hyperintensities (WMHs) in cerebral small vessel disease (CSVD), via Monte Carlo simulations of magnetic susceptibility imaging and a novel imaging marker called the Expected Iron Coefficient (EIC).

**Methods:** A synthetic pathological model of a different number of impenetrable spheres at random locations was employed to represent pathological iron deposition. The diffusion process was simulated with a Monte Carlo method with adjustable parameters to manipulate sphere size, distribution, and extracellular properties. Quantitative susceptibility mapping (QSM) was performed in a clinical dataset to study CSVD to derive and evaluate QSM, R2*, the iron microenvironment coefficient (IMC), and EIC in the presence of WMHs.

**Results:** The simulations show that QSM signals increase in the presence of increased tissue iron, confirming that the EIC increases with pathology. Clinical results demonstrate that while QSM, R2*, and the IMC do not show differences in brain iron, the EIC does in the context of CSVD.

**Conclusion:** The EIC is more sensitive to subtle changes in brain iron deposition caused by pathology, even when QSM, R2*, and the IMC do not.

## 1 INTRODUCTION

White matter hyperintensities (WMHs) are white matter lesions that appear bright on fluid attenuation inversion recovery (FLAIR) MRI, accompany normal aging, and are signs of cerebral small vessel disease (CSVD). WMHs may be caused by sustained oxidative stress throughout the brain, which can be caused by chronic inflammation [1]. Numerous pathologies demonstrate increased brain iron load in the presence of oxidative stress [2], and some of the deep gray matter structures tend to display excessive iron load [3]. However, the exact link between oxidative stress and excessive iron deposition with free radicals in the basal ganglia is not well understood. For example, there are conflicting reports that WMHs lead to accelerated iron deposition throughout the brain, in particular in the following deep gray matter (DGM) structures: caudate (Cau), globus pallidus (GPal), and putamen (Put) [4, 5].

Quantitative susceptibility mapping (QSM) and R2* mapping are among the most popular MRI markers for detecting individual and group changes in brain iron [6, 7]. Non-heme iron deposition in the basal ganglia presents as sub-cellular (≤ 10*µm*, microscopic) and cellular (10*µm* < 100*µm*, mesoscopic) magnetic field inhomogeneities, which manifest in transverse phase decay on gradient echo (GRE) magnitude images [8]. QSM is a macroscopic (≥ 100*µm*) marker of magnetic susceptibility and thus has been shown to reflect brain iron concentration within a voxel, while R2* is sensitive to differences in iron distribution at the mesoscopic scale. Both QSM and R2* demonstrate linear relation-ships with iron in the brain [9, 10], providing a foundation for the simultaneous analysis of QSM and R2* by means of the iron microenvironment coefficient (IMC), introduced and described in Taege et al. [8]. The IMC exploits the linear relationships between magnetic susceptibility and R2* with non-heme iron concentration in structures where iron distribution does not vary systematically throughout a region of interest (ROI), namely the DGM structures. The IMC provides insights into the alteration of DGM tissue as a result of pathology leading to changes in iron deposition and concentration.

While QSM, R2*, and the IMC are all accepted biomarkers relevant to assessing brain iron in tissues, for certain pathologies, standard ROI-based analyses are not sufficient to assess widespread pathological changes in brain iron. For example, heterogeneous findings about iron deposition as a result of mild WMH burden imply that none of the accepted iron-related biomarkers provide definitive insight to more fully understand the role of WMH induced iron deposition [4]. Recent results from our group shows that while QSM is correlated with WMH volume, the standard ROI analyses do not show statistical differences in iron deposition in a cohort demonstrating HIV-associated CSVD with mild WMH burden [11]. Thus, we introduce the Expected Iron Coefficient (EIC) as a novel imaging biomarker, described in Section 2 and use it to evaluate pathological changes in iron deposition across the entire brain in the context of mild WMH burden. By taking advantage of multiple ROIs across the entire brain, we extend the standard mesoscopic and macroscopic analyses to an even larger scale, which may provide better sensitivity in heterogeneous environments to neuropathological patterns of iron deposition.

Specifically, in this work, we evaluated three common iron related imaging biomarkers derived from QSM to assess iron-related tissue changes in the presence of mild WMH burden: QSM, R2*, and the IMC. We further proposed the EIC as a novel population-based imaging biomarker to explore group differences in whole brain magnetic susceptibility to provide insights into increased iron deposition due to pathology. We described the EIC, demonstrated its usefulness by Monte Carlo (MC) simulations, and validated its clinical efficacy using data from a study about HIV-associated CSVD.

## 2 THEORY

### 2.1 Age dependence of tissue iron deposition and magnetic susceptibility

In healthy aging, iron deposition occurs throughout the brain [12, 13, 14], in some regions more than others [3]. Given the unique properties of the brain (e.g., to metabolize, store, and transport iron across the blood brain barrier (BBB)), it may be possible to associate global changes in iron deposition with aging and neuropathology. Hallgren and Sourander [15] experimentally derived equations for calculating iron deposition as a function of age in a typical adult population by posthumously taking tissue samples from regions across the brain of healthy individuals and quantifying the amount of non-heme iron in each tissue.

In this paper, we refer to the natural deposition of iron over time as the expected iron deposition (EID) within an ROI. We assume that these EID formulas represent the amount of iron expected to be present in an individual at a given age without any additional iron metabolizing effects, such as pathology. The EIDs are calculated for each ROI as in Hallgren and Sourander [15] by:

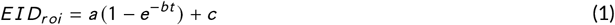

where a, b, and c are experimentally determined region specific constants, t is age in years, and *EID*_*roi*_ is in mg non-heme iron per 100 g fresh weight. In this paper we restrict our study to only those regions with established EID formulas, namely the Cau, Put, GPal, frontal gray matter (FGM), parietal gray matter (PGM), and temporal gray matter (TGM). For single ROI-based analyses, we additionally consider the thalamus (Tha). Table 1 provides values for the region specific constants in Eq. (1).

**TABLE 1.**
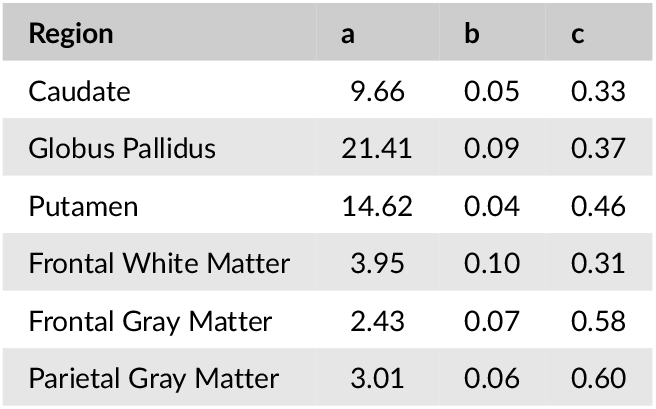
Expected Iron Deposition Coefficients. Regional EID Coefficients given by the least squares regression formulas presented in Hallgren and Sourander [15].

### 2.2 Introduction of the Expected Iron Coefficient (EIC)

We propose a novel group level imaging marker derived from QSM and EID. It is well known that QSM derived susceptibility and non-heme iron concentration are linearly proportional [10]. We hypothesized that this proportionality coefficient can be different between groups of subjects exhibiting different neuropathologies, thus providing a more sensitive marker of subtle iron deposition resulting from pathology. That is, given a cohort of healthy individuals, we would find a group-wise slope between QSM and EID equal to *α*, whereas the slope for a group of subjects impacted by a disease that changes iron deposition patterns across the brain (e.g., mulitple sclerosis (MS), Parkinson’s disease (PD), Alzheimer’s disease (AD), CSVD) would equal *α* ± *ε*, where *ε* indicates a relative increase in iron concentration compared to the healthy group.

These deviations from iron deposition expected with healthy aging lead to a novel group-level imaging biomarker called the expected iron coefficient (EIC). The EIC can be summarized by the following formula:

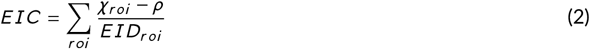

where *χ* is the magnetic susceptibility as measured by QSM, *EIC* is the expected iron coefficient for a given population of subjects, *EID* is the expected iron deposition as calculated by formula (1), and *ρ* is a linear combination of confounding variables that account for additional sources of non-heme iron. Note that *χ* and the EID are not limited to specific ROIs and cannot be used for one single ROI.

#### 2.2.1 Experimental determination of the EIC

The EIC described by formula (2) does not account for additional confounding effects such as age, gender, cardiovascular risk score, and comorbidities that have known effects with iron concentration. Given a group of subjects, we extend our definition of the EIC to include these additional variables. We build a multivariate regression model for *χ* and EID for each subject within a population:

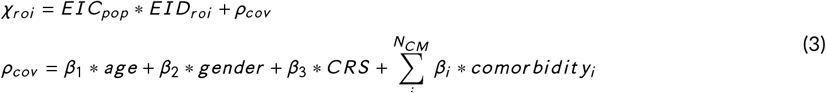

where *EIC*_*pop*_ is the EIC for a given population of subjects, all chosen ROIs are included, and *ρ*_*cov*_ is a linear combina-tion of other variables, including age, gender, cardiovascular risk, and *N*_*C M*_ comorbidities. Therefore, we obtain one EIC for each population in question using susceptibility values in every region chosen for the model (i.e., the Cau, Put, GPal, FGM, PGM, and TGM in this case).

#### 2.2.2 Properties and Limitations of the EIC

Properties of the EIC include:

- **The EIC reflects pathology**. In the derivation of the EIC, we control for age, implying that the EIC is agnostic to aging. Therefore, the EIC is solely reflective of pathological iron-related changes in each study population independent of age.
- **The EIC uses increased SNR by combining multiple regions of interest across the brain**. While some measures are unable to detect iron-related changes in pathologies with subtle iron changes within single ROIs, the EIC uses information from multiple ROIs to reveal pathological effects.
- **The EIC is reflective of brain iron changes**. Some diseases demonstrate localized iron changes (e.g., Huntington’s disease (HD)), while others have a more diffuse effect (e.g., MS, AD, and PD). Since the EIC uses information from multiple ROIs, it is more likely to detect deviations in normal iron deposition resulting from diffuse, whole-brain conditions.

Due to the nature of noninvasive MRI, we cannot be sure that a higher EIC value corresponds to actual increased iron deposition without post-mortem clinical validation. We currently do not have the acquisition sensitivity to image at the cellular and sub-cellular levels. A higher EIC compared to controls implies that there is an increase in average susceptibility from typical aging patterns, not necessarily only non-heme iron concentration. However, since non-heme iron has strong ferromagnetic properties, an increase in susceptibility is likely driven by an increase in iron.

## 3 METHODS

### 3.1 Monte Carlo Simulations

MC diffusion simulations are commonly used to understand some of the underlying biological signals detected with MR pulse sequences [16]. Recent studies have used proton diffusion models to evaluate changes in QSM [17, 18]. The mechanics of the MC simulations used here were employed in Lin et al. [19] and modified to incorporate field inhomogeneities due to magnetic susceptibility sources in the case of a gradient echo acquisition. The following is an overview of the simulation:

1. Generate a random distribution of impenetrable spheres representing varying amounts of iron concentration.

2. Distribute particles (walkers) randomly throughout the volume.

3. Compute the magnetic field at each particle location as a superposition of the field generated from the spheres.

4. Advance the phase of each particle according to its local field.

5. Advance the particles in an MC in an arbitrary direction.

6. If a particle intersects the boundary of a sphere, apply appropriate boundary conditions.

Histological environments in DGM structures are composed of gray matter, white matter, vasculature, and additional cellular sources (e.g., non-heme iron) that lead to magnetic field inhomogeneities [20]. However, due to the paramagnetic nature of non-heme iron to dominate field inhomogeneities in these regions, a simple tissue model consisting only of *N*_*sph*_ randomly distributed spheres of radius *R*_*sph*_ inside a voxel that was 1 *mm*^3^ in volume was implemented to model iron deposition. Additional inhomogeneities were accounted for in the diffusivity parameter. Particles were free to diffuse outside of the cells according to a diffusivity constant *D*_*e*_, reflective of the average diffusivity properties within the DGM structures [21]. This simple model to study magnetic susceptibility has been chosen before in various capacities [22, 23, 24, 25]. *N*_*w*_ = 10,000 walkers were initialized randomly throughout the volume, limited by the computational resources needed for higher values of *N*_*w*_.

A spherical susceptibility source adds the following shift to the magnetic field:

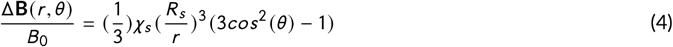

where *B*_0_ = 3*T* is the magnetic field, *θ* is the azimuthal angle with respect to the magnetic field orientation (z-direction), and *χ*_*s*_ and *R*_*s*_ are the magnetic susceptibility and radius of the *s* th sphere. To calculate the total shift in the magnetic field due to all susceptibility sources, a linear superposition of fields was calculated at each location in the volume. Thus, as the walkers diffused, they accrued phase at each TimeStep based on the local field:

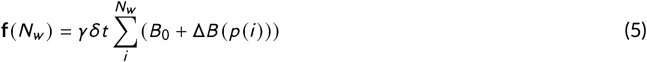

where **f**(*N*_*w*_) is the total phase accrual of all *N*_*w*_ walkers, *p* (*i*) is the position of the *i* th walker, *γ* is the gyromagnetic ratio of the walkers (*γ* = 42.58*M H z* /*T* for Hydrogen), and *δt* is the simulation TimeStep duration. Then, the simulated signal *S* as a function of echo time (TE) of the gradient echo sequence is calculated by

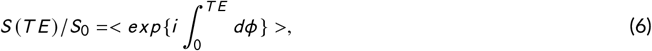

where *dϕ* = 2*π***f**(**r**(*t*))*d t* and < · > indicates an average over all walkers.

To simulate the dynamic diffusive motion of the walkers, a TimeStep duration of *δt* = 0.01*ms* was chosen. Therefore, each walker traveled a distance 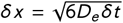 in a randomly chosen direction per TimeStep. We assumed that the water can move freely outside of the iron particles. Within a TimeStep, if a walker collided with an iron sphere, then we assumed a very small transmission probability (i.e., *P* = *e*^−8^) of a water molecule permeating through the cellular membrane. Otherwise, we assumed rigid-body elastic collisions, where the walker was reflected according to the normal vector of the surface at the point of collision. Finally, we assumed periodic boundary conditions, i.e., if a walker exits the volume, then it reenters on the opposite side.

#### 3.1.1 Model of Pathology

We made some assumptions about the neuropathology of iron deposition in the basal ganglia. In order to maintain consistency with the regions with expected iron as calculated in Hallgren and Sourander [15], we restricted our attention to simulate the environments of the Cau, Put, and GPal, only. In general, pathological iron accumulation can be modeled as age-dependent sigmoid functions defined by three parameters in each region. In the present study, we assumed that the rate of increase in iron concentration in the presence of pathology is proportional to the amount of iron expected to be in the region without pathology. More specifically, as simulated QSM is a function of total susceptibility in the environment, to model pathology, we increased the radii of each iron sphere from 20*µm* to 100*µm* to remain consistent with mesoscopic non-heme iron concentrations and distributions [8, 26]. Increasing the radii of the spheres also kept computational resources to a minimum, as opposed to increasing the number of spheres. The simulated signal was calculated by Eq. (6). This model of pathology implies that iron deposition is deposited in clusters, rather than in isotropic distributions. Varying the total susceptibility by increasing spherical radii does not violate physical assumptions about deposition patterns from diffuse oxidative stress, nor does it change the synthesized macroscopic susceptibility signal.

### 3.1 Experimental Validation

#### 3.2.1 Data and Image Processing

In an ongoing study approved by University of Rochester’s Research Subjects Review Board (RSRB), 217 subjects were selected and evaluated to study the effects of CSVD in HIV patients [27]. All imaging was conducted on a research dedicated 3T Siemens PrismaFit scanner (Erlangen, Germany) with a 64-channel head coil. The protocol includes high-resolution T1 MPRAGE images (TI=950ms, TE/TR=3.87ms/1,620ms, 1mm isotropic resolution). QSM data was acquired with a 3D multiple echo GRE sequence (8-echo bi-polar readout train, TE of 1st echo=5.43ms, echo spacing=5.5ms, TR=48ms, readout bandwidth=930Hz/pixel, voxel size=0.9×0.9×2.0 mm^3^, acquisition matrix = 256×256×64). QSM and R2* images were calculated using the Morphology Enabled Dipole Inversion with zero-tissue referencing (MEDI+0) toolbox, using default normalization settings [28, 29]. T1-weighted images were used for coregistration. ROIs were selected based on the available formulas in Hallgren and Sourander [15] and applied as masks in native QSM space in FSL [30]. FLAIR images were collected and CSVD burden was assessed via deep and periventricular Fazekas scores by our team’s radiologist [31].

We assessed cardiovascular risk via the Reynold’s risk score [32] as an additional clinical covariate. Next, we separated the remaining subjects based on two conditions: the presence of HIV-infection and the presence of WMH burden via deep and periventricular Fazekas scores of one or higher [31], consistent with the total CSVD burden score [33]. Subjects with deep and periventricular WMH scores of zero and one were removed due to very small WMH burden. This resulted in four cohorts described in table 2: Healthy controls (HIV-CSVD-), controls with WMH burden (HIV-CSVD+), subjects with HIV-infection (HIV+CSVD-), and subjects with HIV-infection and WMH burden (HIV+CSVD+).

**TABLE 2.** Study Demographic Information. Where appropriate values are given as mean (SD).

| Cohorts | HIV-CSVD- | HIV-CSVD+ | HIV+CSVD- | HIV+CSVD+ |
| --- | --- | --- | --- | --- |
| Number of Subjects | 28 | 16 | 22 | 34 |
| Gender (male:female) | 17:11 | 8:8 | 15:7 | 22:12 |
| Age (years) | 44 (15.6) | 61 (11) | 44 (9.9) | 58 (6.0) |
| Reynold's Risk Score | 4.0 (5.1) | 7.2 (8.5) | 2.6 (2.1) | 9.9 (7.7) |

#### 3.2.2 Biomarker Experiments

##### ROI-based Markers

QSM, R2*, and IMC group differences were calculated by four-way ANCOVA analysis followed by post-hoc testing via Welch’s t-tests. For each DGM ROI for each subject, the IMC was calculated as follows: (1) a Pearson correlation was performed per voxel between QSM and R2* maps; (2) A total weighted least squares linear regression was performed between QSM and R2* values per voxel. This regression coefficient (*κ*_*other*_) is the IMC defined by formula (**??**).

##### EIC Marker

We explored the relationship between absolute average susceptibility and expected iron deposition across various brain regions. Here we regressed the average QSM values against the EIDs in the frontal, parietal, and temporal cortices, Cau, Put, and GPal, controlling for age and Reynold’s Risk Score in order to use the regression coefficient as a biomarker for WMH burden. We considered the four cohorts described above and perform ANCOVA analyses to determine significant differences between EICs.

The multivariate regression models included in this analysis are described below. The full model (Eq. (2)) is written explicitly for a fixed-effects linear model:

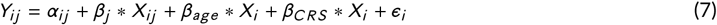

where *β*_*j*_ is the EIC, *α*_*j*_ is the intercept, *j* = 1, 2, 3, 4 corresponds to each cohort, i represents each observation (subject), and age and Reynold’s Risk score are considered as covariates. Note that since we use six regions of interest, there are 6 * *N*_*sub*_ observations, where *N*_*sub*_ is the number of subjects. The clinical covariates were included 6 times for each subject. The full regression model was compared to a restricted model where *β* = *β*_*j*_ (i.e., all slopes are equal):

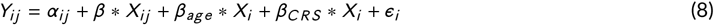

An F-test between the two models was performed. Post-hoc group t-testing was also performed to directly compare whether the diseased cohorts were different from the HIV-CSVD-group.

## 4 RESULTS

### 4.1 Simulations

To validate the performance of the simulations, the signal was calculated according to equation (6) as a function of TE (time of first echo = 5.43ms, equal echo spacing = 5.5ms, TR = 48ms) for one preset geometry and run 20 times to estimate signal variability. Mean signals are plotted using different numbers of random walkers (Figure 1). As the number of walkers increased, the exponential decay expected from T2* relaxation became more consistent and less variable. For all further experiments, *N*_*w*_ = 10, 000 walkers were chosen due to computational resource limitations (memory storage and computational times exceeded hardware limits for *N*_*w*_ > 20, 000). The signal variability (standard deviation over twenty simulations) at each TE was less that 0.001 for *N*_*w*_ = 10, 000. All simulations were performed in MATLAB version 2019b (MathWorks, Natick, MA) on an Intel CPU (2.70 GHz).

**FIGURE 1.**
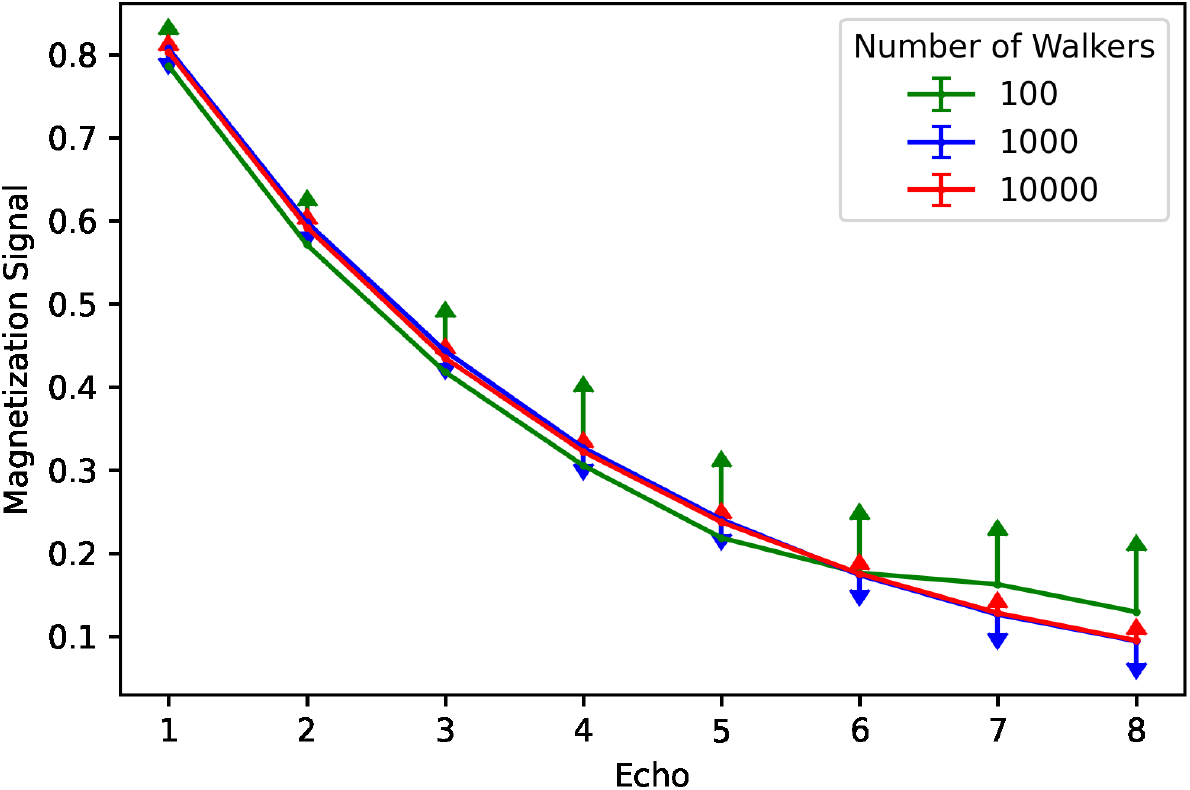
Simulation signal validation. The magnetization signal is plotted for each echo (constant echo spacing) for different numbers of walkers in the simulation. Error bars are displayed based on 20 simulations run for each number of walkers. The signal becomes less variable and more consistent with a typical T2* decay with more walkers.

Additional validation simulations were performed to confirm that the signal reflects the underlying pathology. As QSM is reconstructed by taking advantage of a linear phase accrual as protons diffuse throughout an inhomogeneous local magnetic field, we confirmed that the rate of phase accrual as a function of time increases as total susceptibility within the environment increased. This relationship held consistent when holding either the number of spheres or total susceptibility constant. Therefore, the Monte Carlo simulations provide information about the total iron deposition in the local environments, namely that the rate of phase accrual reflects changes in iron deposition and the calculated signal reflects T2* decay (1/R2*), indicative of changes in iron distribution.

From the simulated pathology, for a given number of iron spheres, the radius of each sphere was increased from 20*µm* to 100*µm* in order to represent clusters of increased iron deposition, where larger clusters correspond to higher pathological iron deposition. Figure 2 shows the rate of phase accrual as a function of total susceptibility, by region of interest (number of spheres). The different number of spheres represent the susceptibility levels of the Cau (*N*_*sph*_ = 5), GPal (*N*_*sph*_ = 15), and Put (*N*_*sph*_ = 10). The colors indicate the radii of each of the spheres in the simulated environment. Normal aging (dark blue) is considered to have radii equal to 20*µm* and increasing radius (up to 100*µm*) indicates increased iron deposition as a result of pathology (dark red). The increased radii of the spheres represent iron deposition that is concentrated to specific clusters within the DGM tissues. However, given that QSM measurements are agnostic to iron distribution, the locations of the increased iron load does not influence the macroscopic susceptibility measurements. A relatively small number of spheres was chosen for each ROI due to computational demands.

**FIGURE 2.**
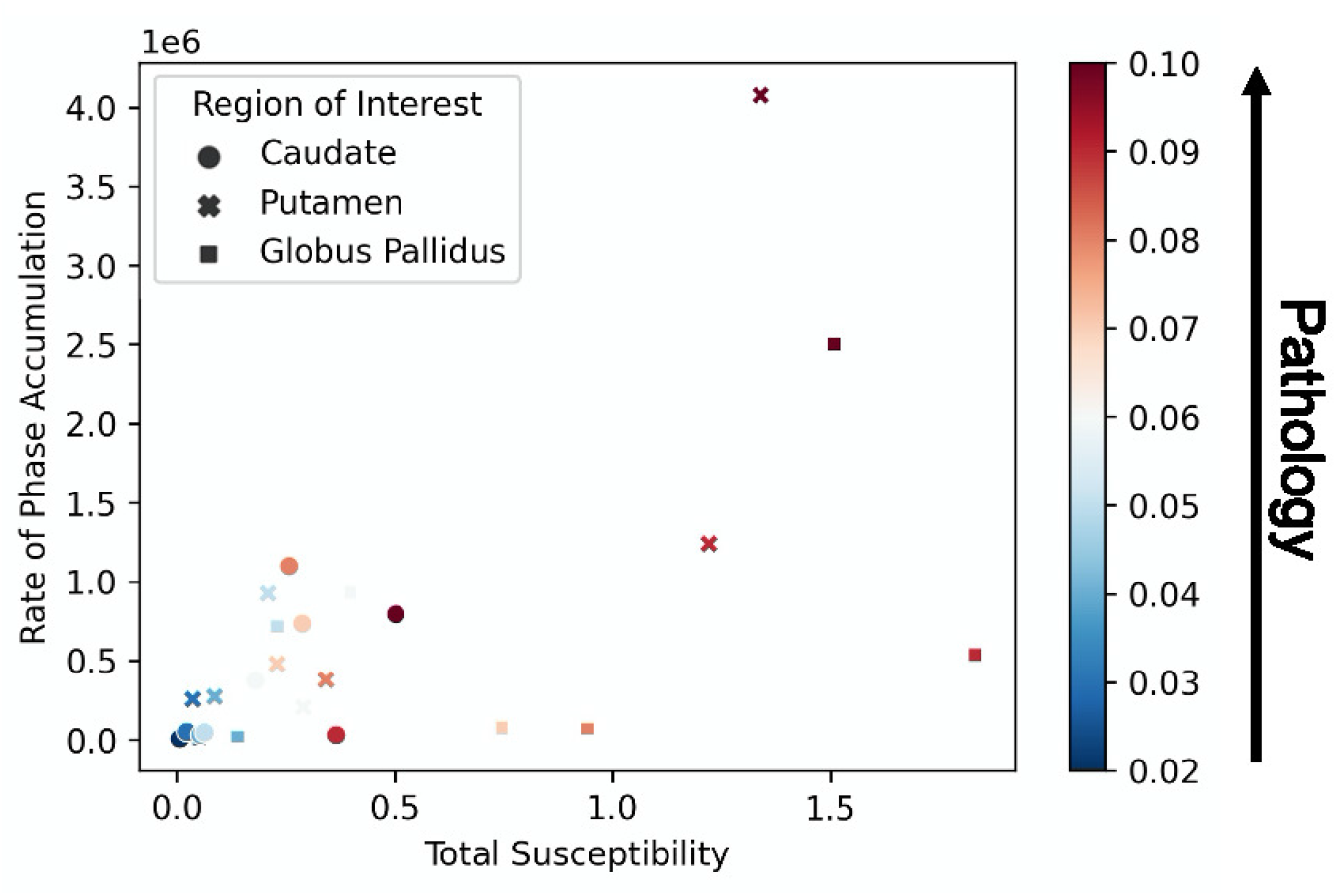
Pathology Simulation Results. The rate of cumulative phase accumulation is plotted against the total susceptibility. 5, 10, and 15 spheres were used to simulate the Caudate, Putamen, and Globus Pallidus, respectively. Pathology was increased by increasing the radius of each sphere in every region. The rate of phase accumulation and total susceptibility increase as a result of increased pathology.

### 4.2 Clinical Experiments

Figure 3 shows the regions of interest included in these analyses and example R2* and QSM maps from one healthy subject in standard space. Figure 4 shows the mean and standard error of the QSM, R2*, and IMC markers in the Cau, GPal, Put, and Tha. There were no differences found in the ROI based analyses for QSM, R2*, nor the IMC via ANCOVA analyses and post-hoc t-tests between groups.

**FIGURE 3.**
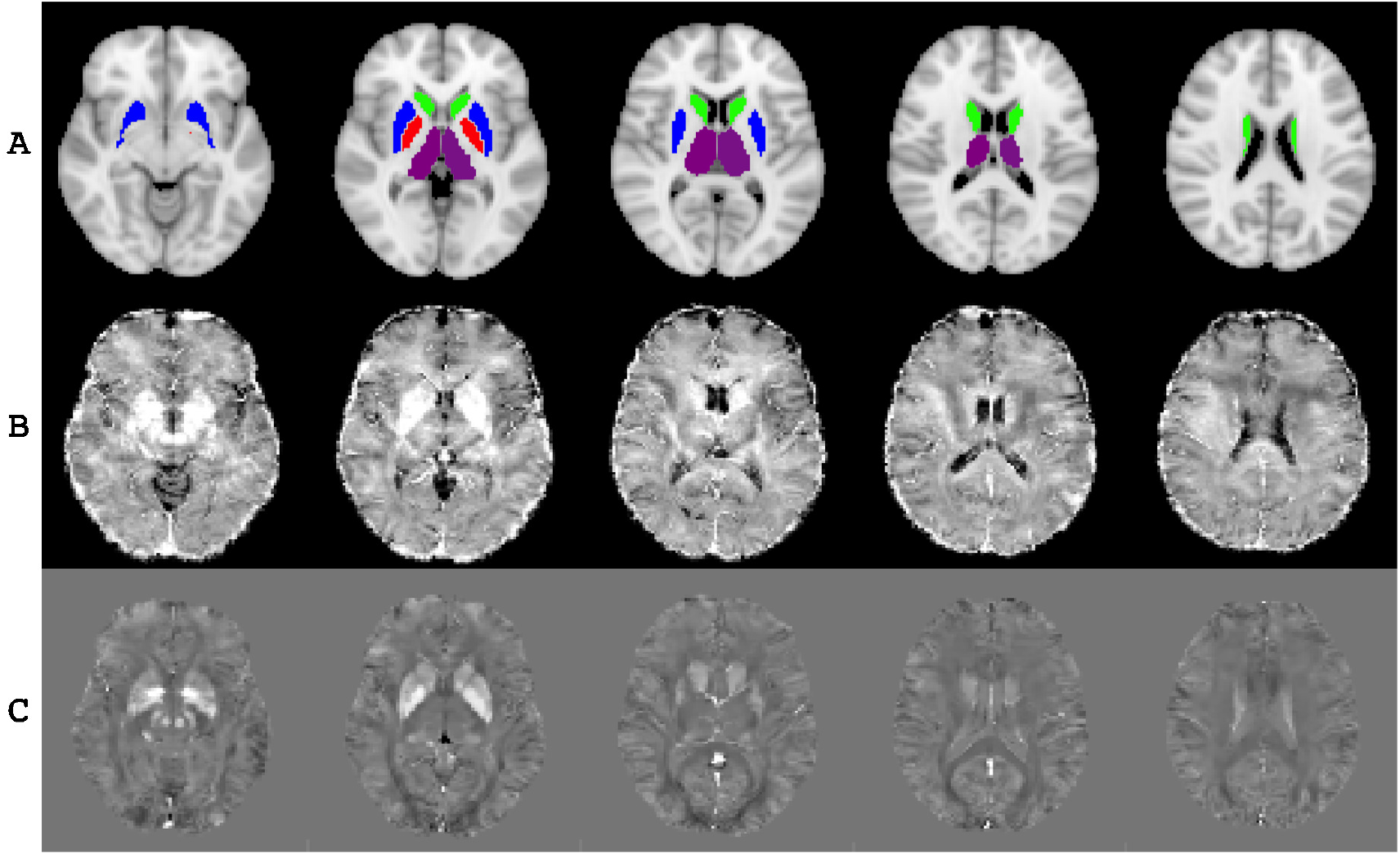
A. Regions of interest selected for consideration: Cau, green; GPal, red; Put, blue; and Tha, purple. Example B. R2* and C. QSM maps of one healthy subject in standard space.

**FIGURE 4.**
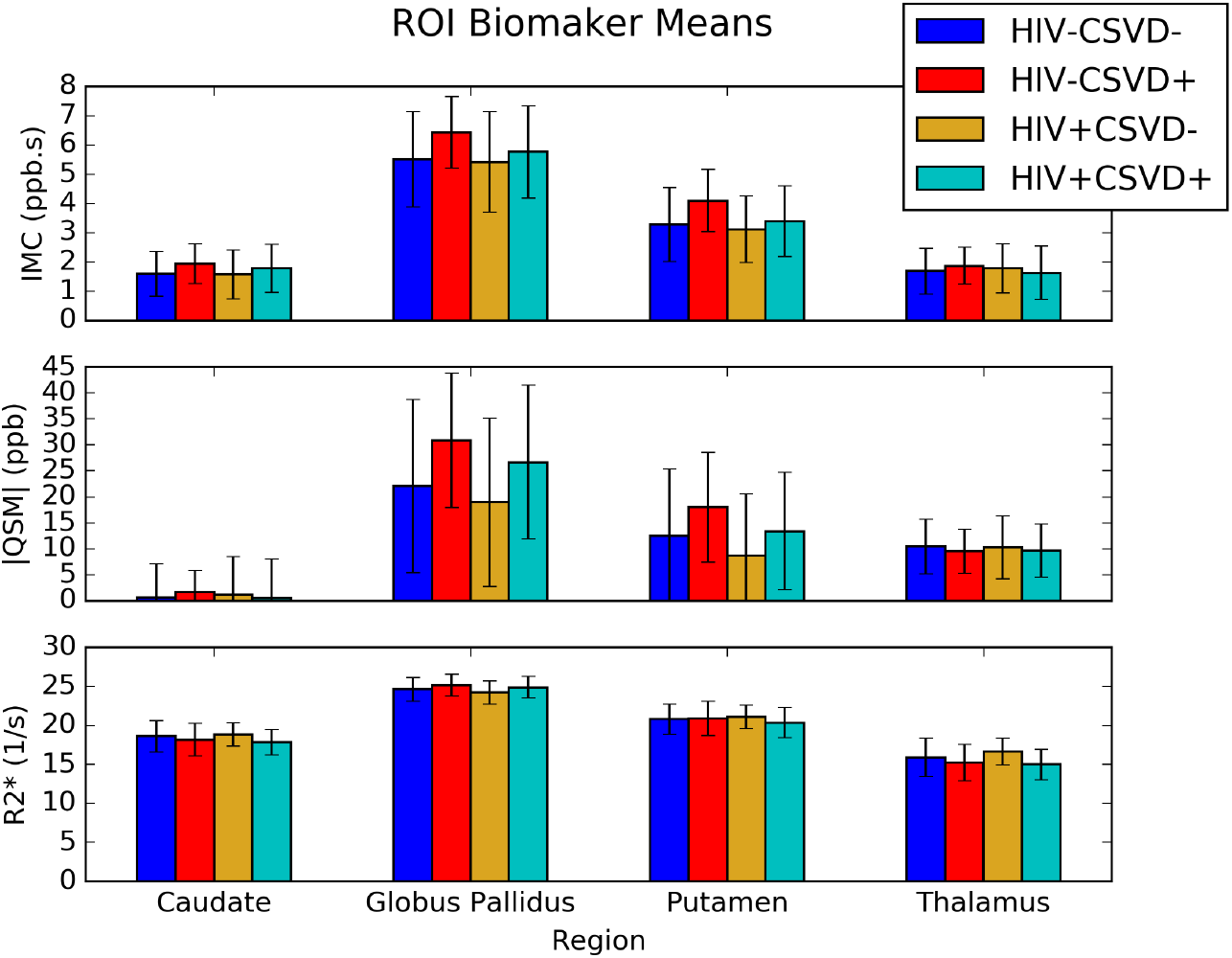
Barplots of region of interest biomarker (means with standard deviation) for the iron microenvironment coefficient (IMC), units ppb s, absolute QSM, units ppb (parts per billion), and R2*, units *s* ^−1^. None of the biomarkers show any statistical differences between groups.

Figure 5 shows a scatter-plot of susceptibility values against expected iron deposition for all regions of interest. Regression lines are plotted for each cohort. The slope of each regression line represents the EIC for the population in question. This plot shows that all four cohorts have linear relationships with expected iron deposition. Further, the HIV-CSVD-and HIV-CSVD+ EICs are significantly different.

**FIGURE 5.**
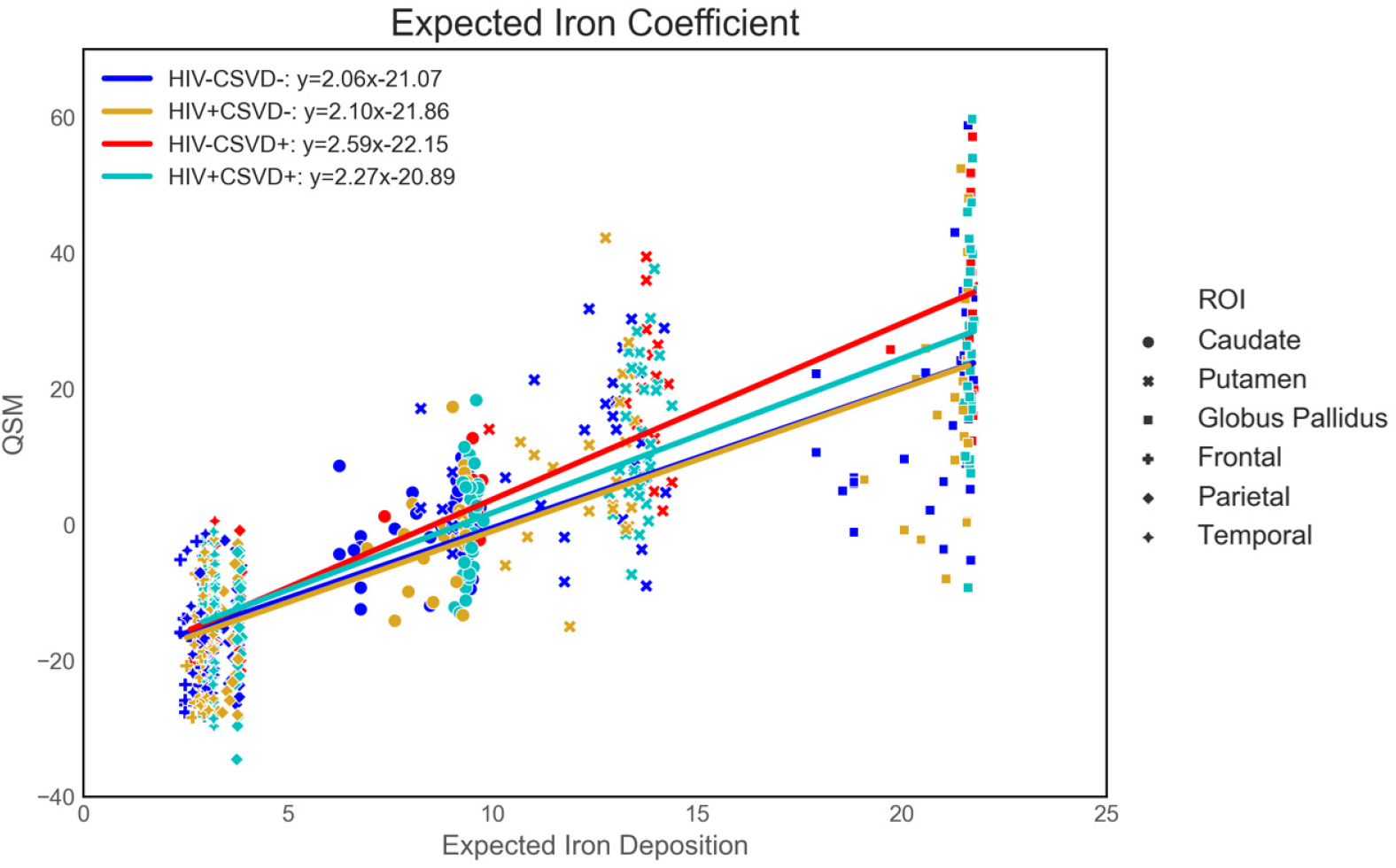
Scatter-plot of QSM vs. EID separated by cohorts for six regions of interest across the brain. Regression lines indicate linear fit and equations are displayed for each cohort. The EIC is represented by the slope of the regression lines for each group. There is a significant difference between the EICs of the HIV-CSVD-and HIV-CSVD+ groups.

Additional regression results for each population are given in Table 3. The HIV-CSVD-group (healthy controls) was selected as the reference group. The EIC_*j*_ and Int_*j*_ for the cohorts are relative to EIC_*H IV* −*C SV D*−_ and Int_*H IV* −*C SV D*−_, respectively. The significance of EIC_*H IV* −*C SV D*+_ shows that the EIC difference between the HIV-CSVD-and HIV-CSVD+ cohorts is significant and independent of clinical covariates. It is also worth noting that in both models, Reynold’s risk score is significant.

**TABLE 3.** Full and Restricted Model regression results, where the full and restricted models are given by Eqs. (7) and (8), respectively. CRS, cardiovascular risk score.

| Full Model |  |  | Restricted Model |  |  |
| --- | --- | --- | --- | --- | --- |
| Variable | Coefficient | P-value | Variable | Coefficient | P-value |
| $EIC_{HIV-CSVD-}$ | 2.04 | <0.001 | EIC | 2.20 | <0.001 |
| $Int_{HIV-CSVD-}$ | -24.18 | <0.001 | $Int_{HIV-CSVD-}$ | -25.33 | <0.001 |
| Age | 0.05 | 0.192 | Age | 0.04 | 0.237 |
| CRS | 0.31 | <0.001 | CRS | 0.07 | <0.001 |
| $EIC_{HIV-CSVD+}$ | 0.46 | 0.011 | $Int_{HIV-CSVD+}$ | 1.43 | 0.296 |
| $Int_{HIV-CSVD+}$ | -2.73 | 0.198 | $Int_{HIV-CSVD-}$ | -0.41 | 0.707 |
| $EIC_{HIV-CSVD-}$ | 0.06 | 0.725 | $Int_{HIV+CSVD+}$ | -0.24 | 0.827 |
| $Int_{HIV-CSVD-}$ | -0.89 | 0.621 | | | |
| $EIC_{HIV+CSVD+}$ | 0.22 | 0.139 | | | |
| $Int_{HIV+CSVD+}$ | -2.15 | 0.211 | | | |

An F-test comparing the two models showed equal variance between overall models. T-tests of EICs between cohorts showed that the HIV-CSVD-cohort was significantly different from the HIV-CSVD+ cohort (p_*F DR*_ =0.000072). No other EIC comparisons showed significant differences. Regression models were also performed including the three subcortical regions only, excluding tissue from the cortices. Results were similar with a slightly lower corrected p-value for T-tests between EICs of the HIV-CSVD-and HIV-CSVD+ groups (p_*F DR*_ =0.000052).

## 5 DISCUSSION

This study used 3D Monte Carlo simulations of a multiple-echo GRE sequence and clinical data to evaluate changes due to iron deposition via magnetic susceptibility in the DGM structures caused by chronic neuropathology. This study introduced a population-based imaging marker derived from QSM, called the Expected Iron Coefficient (EIC), which is reflective of pathological iron deposition across a population of subjects. Further, we demonstrated that the EIC has a clinical advantage over ROI-based brain iron markers in neuropathologies that cause subtle increases to iron deposition in the DGM brain structures most sensitive to iron accumulation.

Our simulation results showed that using a multiple-echo GRE pulse sequence, phase accrual of all protons in a simulated environment (i.e., voxel) is linear and driven by local magnetic field inhomogeneities (e.g., susceptibility sources). These results are consistent with what has been previously shown with regard to QSM and R2* mapping, namely that QSM measurements reflect the macroscopic nature of the susceptibility within a voxel while R2* provides information about the within-voxel susceptibility distribution [8].

From our pathology simulation results, we showed that as particular ROIs, denoted by different numbers of spheres of deposited iron, increase in total susceptibility, the rate of cumulative phase accrual increases. By simulating environments with different base levels of iron accumulation (20*µm*, healthy) and increasing radii (up to 100*µm*, pathology) within the mesoscopic spatial range, we found that QSM and R2* signals increase with pathology. These simulations also suggest that iron accumulation resulting from neuropathology is in addition of typical aging. The results in Figure 2 show increasing pathology for a single individual for three separate ROIs. In every region, QSM increases as a function of pathology at any age. Therefore, when many subjects are involved in a clinical dataset, the EIC is a marker reflecting increased iron deposition as a direct result of pathology.

In our clinical population, we considered the effects of HIV-infection and WMH burden, indicating CSVD. CSVD is more prevalent in the presence of HIV [34] and chronic inflammation in the white matter may lead to oxidative stress in the brain and thus increased iron deposition in the DGM brain structures compared to age-matched controls [35]. By comparing three iron-derived imaging biomarkers (i.e., QSM, R2*, and IMC) between the four groups stratified by HIV and CSVD statuses (Figure 4), we found trends that suggest alterations in iron deposition and distribution in the DGM, despite lacking statistical significance in single ROI-based tests. For example, within both the HIV infected and uninfected cohorts, subjects with CSVD showed increased QSM, R2*, and IMC in nearly every ROI. Alternatively, subjects with HIV tended to display slightly lower QSM and IMC levels than their respective control groups. Neither of these trends can be used to draw definitive conclusions about the nature of pathological iron deposition in HIV nor CSVD.

However, applying the EIC to our four groups, we did find one strong and consistent finding. The EIC of the HIV-CSVD+ group was significantly higher than both groups without signs of CSVD. Pooling all subjects based on HIV status showed that the CSVD+ group had a significantly higher EIC than the CSVD-group, irrespective of HIV infection. Further, the EICs of the HIV+ and HIV-groups without CSVD overlap, suggesting that HIV infection not does influence the EIC. These results imply that WMH burden does lead to increased iron deposition in the DGM and throughout the brain, while HIV infection does not. Future work should be performed to assess the trends of decreased QSM due to HIV, which may be attributable microglial dysfunction in HIV infection [36]. The increase in EIC in the CSVD+ groups indicates that the rate of iron deposition is higher after controlling for age and cardiovascular risk score, both of which are known to be associated with increased QSM [10, 11]. For studies involving low rates of iron deposition in the brain, the EIC provides more information about the changes in tissue susceptibility than the standard ROI-based analyses. This increased sensitivity to detect subtle changes in iron deposition should be extended to other pathologies known to impact brain iron with intermediate stages, such as mild cognitive impairment (MCI) and HD [37, 38].

The work of Hallgren and Sourander [15] has been used to show linearity of QSM and iron deposition in MS [39], PD [40], and healthy aging [9]. However, none of these studies have considered the differences in linearity between diseased groups and healthy controls. The present study explored the differences in this linearity between QSM and iron deposition as a direct result of pathology, providing a novel marker to explain subtle deviations in iron deposition from typical aging. Further, the EIC should become even more apparent in pathologies with larger QSM differences, in particular in populations that do display significantly higher QSM levels using conventional ROI-based analyses, such as MS [39].

We are not aware of any tissue-based Monte Carlo simulation studies that have been used to directly assess the effects of pathological iron deposition in this context. However, previous studies have used simulations to demonstrate that QSM is linearly proportional to the total magnetic susceptibility in a voxel [18] and to explore the relationships between iron distribution and concentration within a simulated environment [8]. Further, a pathological simulation study involving tissue susceptibility was performed to plot trajectories of population values [39], but within-voxel environment interactions were not considered.

One limitation of this study was the tissue model used to simulate the DGM structures. Given the computational complexity to account for the complexity of DGM structure [20], only spherical iron was included as a simulation parameter and the remaining effects were accounted for in the diffusivity coefficient, which was considered to be constant with increasing pathology, as the gross histological makeup of DGM structures is not known to significantly change with respect to mild pathology [41]. Additionally, given the dominating paramagnetic contribution of iron to influence QSM signals, we did not include effects of demyelination nor calcification to minimize computational complexity. Another limitation of the simulations was the computational resources necessary to perform simulations with larger numbers of walkers. We expect that the simulated signals would become more stable with more walkers, as demonstrated in Figure 1, but the time and memory needed to complete the simulations exceeded machine limits with a number of spheres greater than 20 and number of walkers greater than 20,000.

One limitation of our clinical population is a small WMH burden. Many subjects in both the HIV+ and HIV-groups did not qualify to have mild signs of CSVD. As such, many subjects were excluded from the total dataset to maximize the contribution of pathology on QSM levels. A population with higher WMH burden would likely show an even higher EIC and thus rate of iron deposition. Another clinical limitation is the number of ROIs selected to build the EIC regressions. Hallgren and Sourander [15] is the only histological sample of which we are aware that has calculated non-heme iron deposition as a function of aging in a typical adult population. However, as iron in the deep gray matter becomes more well studied as a function of age [42], we expect the EIC to become more stable if additional ROIs are included in the model throughout the brain, because the EIC relies on combing QSM information from many areas across the brain.

## 6 CONCLUSIONS

In this study, we proposed the EIC as a novel imaging marker derived from QSM. The EIC was reflective of deviations in DGM iron deposition compared to age-matched controls and showed that mild WMH burden leads to increased iron deposition in the DGM. We performed Monte Carlo simulation experiments to show that increased iron deposition as a direct result of pathology leads to an increase in tissue susceptibility, confirming that the EIC reflects changes in levels of DGM iron compared to age matched controls. We also applied the EIC to a clinical dataset from a study about HIV-associated CSVD to demonstrate that the EIC provides new information about iron deposition in the DGM associated with the presence of WMHs than is available from standard analyses using QSM, R2* and the IMC. Even though the exact nature of the relation between WMHs and increased iron deposition in the DGM requires further elucidations, we did find that the EIC was associated with mild WMH burden, suggesting higher rates of iron deposition than typical aging.

## Acknowledgements

We would like to acknowledge study participants involved in the clinical dataset; study coordinators Teresa Oh, Jill Guary, Gillian Crysler, and Valerie Kline; Xing Qiu and Will Francis-Consagra for statistical consultations; and the Schifitto imaging laboratory for data collection and technical support including Abrar Faiyez, Alan Finkelstein, Arun Venkataraman, and Yuchuan Zhuang. This work was funded by the NIH grant R01-AG054328, obtained by Giovanni Schifitto and Sanjay Maggirwar.

## Conflict of Interest

The authors have no conflicts of interest to declare.

### Abbreviations

CSVD: cerebral small vessel disease
DGM: deep gray matter
EIC: expected iron coefficient
EID: expected iron deposition
HIV: human immunodeficiency virus
IMC: iron microenvironment coefficient
MRI: magnetic resonance imaging
QSM: quantitative susceptibility mapping
ROI: region of interest
WMH: white matter hyperintensity.

